# GGE: General-purpose deep meta-learning for classification of human transcriptomes with limited data

**DOI:** 10.64898/2026.09.27.754054

**Authors:** Gal Yankovitz, Irit Gat-Viks

**Affiliations:** The Shmunis School of Biomedicine and Cancer Research, George S. Wise Faculty of Life Sciences, Tel Aviv University, Tel Aviv, Israel

**Keywords:** Transcriptome data, deep meta-learning, human foundation model, few-shot classification, MAML, TabNet

## Abstract

Transcriptomic classification is often hindered by the small number of samples relative to the high dimensionality of gene expression data. We introduce General Gene Expression (GGE), a deep meta-learning framework designed to support robust classification in this limited-sample setting. By training across 5,220 distinct biomedical prediction objectives drawn from 1,779 different human datasets, GGE learns a model initialization that captures biological patterns shared across heterogeneous classification tasks. This learned initialization has two key advantages. First, it enables improved performance to new datasets using only a small number of labeled samples. Second, because it is learned across diverse prediction objectives, it can be applied to a broad range of biomedical problems. We show that GGE outperforms established classifiers in data-limited settings across a wide range of applications, including datasets generated using different RNA-seq platforms and preprocessing pipelines. In addition, attention-based analysis identifies recurrent genes that contribute to performance across multiple biological objectives, providing insight into shared determinants of human biological states. Together, these results establish GGE is a general-purpose framework for human transcriptome-based classification in biomedical settings where labeled data are scarce.

## BACKGROUND

Basic biology and biomedicine increasingly rely on genome-wide gene expression profiling to address diverse research and clinical questions. Despite the rapid growth in the number of transcriptomic datasets and the expanding range of their applications, many individual gene-expression studies remain small^1^. This reflects fundamental constraints inherent to biological research: targeted perturbation experiments, assessments of response to therapy, and rare disease studies are difficult to conduct and expensive to recruit at scale. As a result, transcriptomic analysis is commonly performed in a small-n, large-p setting, in which the number of measured genes (p) greatly exceeds the number of samples (n), limiting statistical power and reducing the robustness of discovery.

An intuitive strategy to address limited sample sizes is to integrate information across multiple studies. In practice, however, such integration is often avoided due to concerns about batch effects and cross-study heterogeneity. Instead, many computational workflows follow a meta-analysis strategy in which each dataset is analyzed separately and the results are combined only at a later stage^2^. While this cautious approach can reduce technical bias, it may also limit gains in statistical power available for initial discovery.

Recent advances in *few-shot learning* provide an alternative approach to obtain accurate classification from only a handful of labeled samples. In few-shot learning, a model is pretrained on one or more *source* datasets and then adapted to a *target* dataset with only a small number of labeled samples, instead of learning model parameters from scratch. This paradigm is implemented via transfer learning or meta-learning^3,4^, and multiple studies show that when the source datasets are biologically aligned with the target dataset, pretraining can substantially improve performance in low-sample settings^5–11^. For example, few-shot models pretrained to predict drug response in one tissue can be adapted to predict response of the same drug in another tissue^12,13^, and few-shot models pretrained on several types of cancer can be adapted to an additional type of cancer^8^. However, existing few-shot applications in transcriptomics still rely on carefully chosen source datasets whose relevance to the target problem is determined based on prior knowledge. This creates two limitations. First, these methods are difficult to apply when closely related source and target datasets are not available. Second, they make limited use of the broader public transcriptomic datasets, implicitly assuming that only closely matched conditions can provide useful transferable information.

Growing evidence suggests that the same core transcriptional programs are activated in distinct biological conditions, differing mainly in magnitude or relative composition rather than in their fundamental nature. For example, data from rare monogenic immune diseases have been shown to facilitate interpretations of common complex diseases^14^. Similarly, integration of data from murine infected lung tissue^15^ with data from human blood during chronic low-grade inflammation^16^ improves stratification of sepsis patients^17^. This observation motivates a few-shot learning approach that does not depend on matching between source and target data, but instead exploits the sharing of transcriptional programs across diverse biological conditions.

Here, we developed the General Gene Expression (GGE) model, a meta-learning–based few-shot predictor for human bulk-transcriptome data, implemented with MAML^18^ and TabNet^19^. By pretraining on 1,779 public human RNA-seq datasets spanning diverse cell types, health conditions, and experimental settings, GGE learns features that generalize across biological contexts. The pretrained model can then be adapted to a new classification objective using only a few labeled samples. Accordingly, GGE does not require closely matched source and target cohorts, but instead draws on the breadth of public human transcriptomic data to enable robust prediction in data-limited settings.

Across multiple independent transcriptomic datasets, GGE outperformed alternative classification approaches under limited-sample conditions, with robust results across diverse biomedical contexts, RNA-seq technologies, and preprocessing strategies. Analysis of the model’s attention patterns further revealed genes repeatedly used across distinct prediction tasks, pointing to common axes of human diversity. Taken together, these findings support GGE as a broadly applicable framework for transcriptome-based prediction when only small labeled cohorts are available.

## METHODS

### Overview: A general-purpose few-shot classifier for gene expression data

Our objective is twofold: to enable robust classification in new datasets with limited sample size and to develop a method that can be applied across diverse research objectives without requiring a tailored prior dataset for each new dataset. To address this challenge, we adopt a few-shot meta-learning framework, in which a model is first *pretrained* on a collection of *source tasks* and is then *adapted* to a new *target task* using only a few samples (**Figure 1A**). Here, a *task* is defined by (i) a gene-expression dataset and (ii) a research objective specified by class labels for each sample, such as disease versus healthy or with versus without treatment (**Figure 1A, left**). We refer to the model trained on all source tasks as the *pretrained model*, and to the model after task-specific adaptation as the *adapted model*. For any given task, the adapted model serves as the task-specific classifier. Particularly, we implemented the framework using the Model-Agnostic Meta-Learning (MAML) algorithm^18^, which pretrains across multiple heterogeneous source tasks and therefore the same pretrained model can be adapted well to a wide range of new research objectives. We refer to model pretraining with the MAML algorithm as *meta-training*.

**Figure 1.**
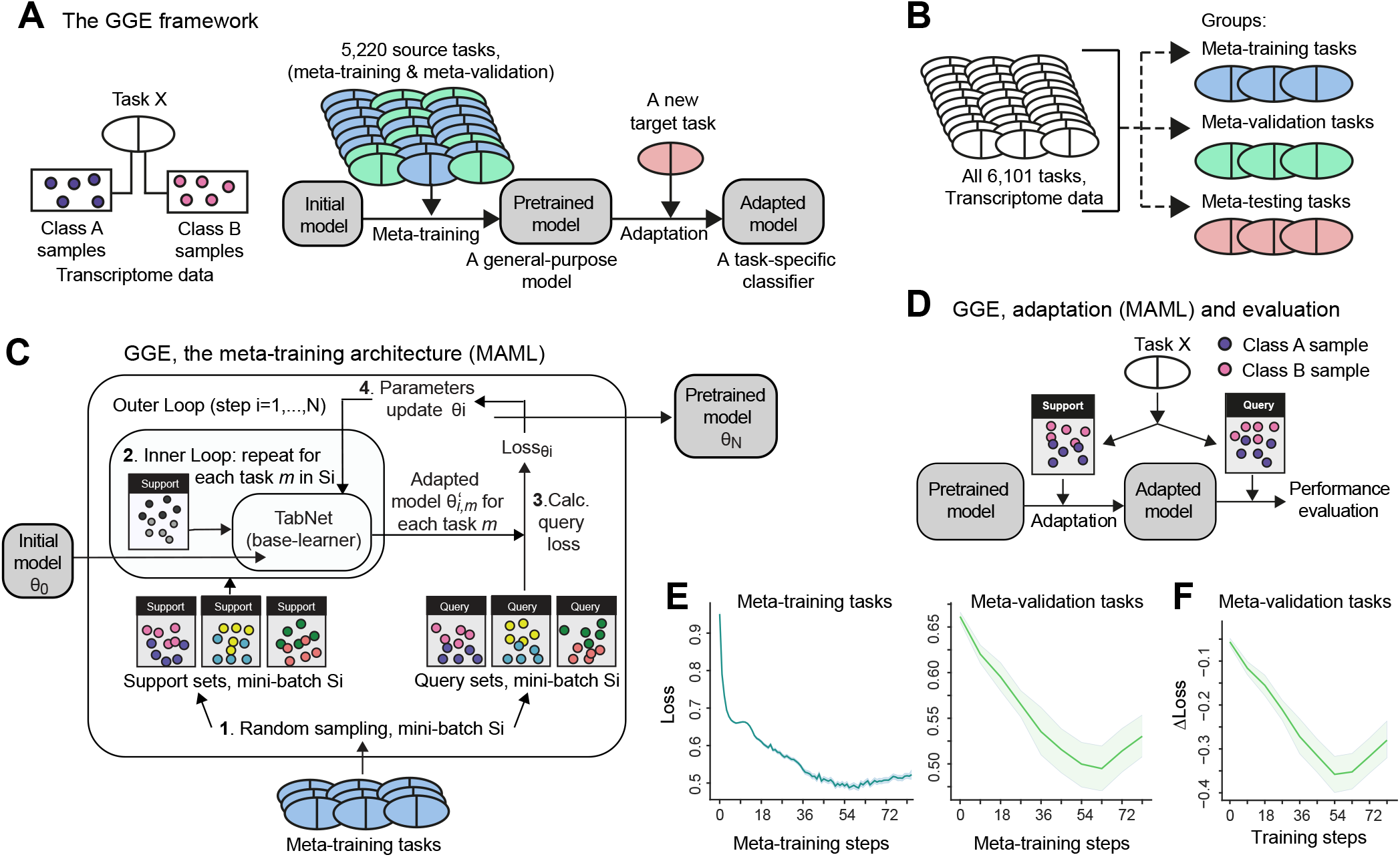
Overview and validity of the GGE framework. (**A**) General-purpose few-shot classification with meta-learning. The GGE model was first meta-trained on a large compendium of source tasks to learn a robust parameter initialization. Next, it can be adapted to a new target task using only a small number of labeled samples. (**B**) Definition of tasks and data splits. The datasets are divided into three non-overlapping splits used for meta-training, hyperparameter tuning (meta-validation tasks), and model evaluation (meta-testing tasks). (**C**,**D**) Schematic illustration of meta-training and adaptation. Within each task, samples from each class are randomly divided into support and query sets, which are used separately during meta-learning and evaluation. (**E**) Performance across meta-training steps, showing progressive improvement on the meta-training tasks (left) and meta-validation tasks (right). (**F**) Benefit of task-specific adaptation. Shown is the Δloss metric that quantifies the improvement on the query set induced by adaptation on the support set. Results reported across the meta-validation tasks.

For this framework, we assembled a large collection of tasks derived from human transcriptomic datasets spanning diverse biological conditions and experimental settings. Tasks were systematically extracted from the Gene Expression Omnibus (GEO) by identifying datasets containing two annotated biological conditions that could serve as class labels (**Methods**). Overall, the collection of tasks used in this study consists of 6,101 tasks from 2,079 distinct transcriptome datasets. The tasks were partitioned at the dataset level into three non-overlapping splits: meta-training tasks (to meta-train the pre-trained model; 1,479 datasets, 4,311 tasks), meta-validation tasks (to tune the hyperparameters of the model; 300 datasets, 909 tasks), and meta-testing tasks (for performance evaluation; 300 datasets, 881 tasks), such that all tasks derived from a given dataset were assigned to a single split (**Figure 1B**). Each task is further partitioned into support and query sets. During meta-training, the support set is used for task-specific parameter updates, whereas the query set is used to update a shared initialization across tasks (**Figure 1C**). To assess performance, the model is first adapted on the support set of each task and then evaluated on its query set (**Figure 1D**). Unless stated otherwise, all reported performance metrics are computed on the meta-testing tasks.

We focused on binary (two-way) classification in a 5-shot classifier. MAML is compatible with different choices of the underlying classifier, called “base learner”. For this study, we selected TabNet^19^ as the base learner, given its suitability for tabular data such as genome-wide bulk transcriptome profiles (**Figure 1C, Methods**). We refer to the resulting MAML–TabNet framework, meta-learned under the two-way 5-shot setting with thousands of heterogeneous tasks, as the *General Gene Expression (GGE)* model.

### Architecture of the meta-training framework

MAML involves two components: a ‘base-learner’ and a ‘meta-learner’. The base-learner (here, TabNet^19^) is a model that learns to solve a specific task, and the meta-learner (here, MAML with FOMAML implementation^18^) is an outer-loop process that trains the base-learner across a variety of tasks, with the objective of making it an effective learner on new tasks. The base-learner adapts to the task using the support samples, and the meta-learner updates its parameters based on the base-learner’s performance on the query samples of all tasks. This forces the model to learn how to adapt effectively from limited data.

The MAML architecture includes inner and outer loops^18^. At each meta-training step_*i*_, a subset *S* _*i*_of meta-training tasks (referred to as a *mini-batch*) is sampled, and each task is randomly divided into disjoint support and query sets. *i*) *Inner loop*. A TabNet network is initialized with the meta-parameters from the previous meta-training step, *θ*_*i–1*_, and is subsequently adapted to the support set of each task *m* independently of other tasks in the mini-batch. Adaptation is applied through several gradient steps with a *learning rate α*, yielding task-specific *adapted parameters* 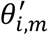(*θ*_0_ at the first step). We use the pytorch-tabnet implementation with variance-preserving Glorot normal initialization^20^. *ii*) *Outer loop*. The outer loop updates the shared meta-parameters based on model performance on the query sets of all tasks in the mini-batch *S*_*i*_. For each task *m* from mini-batch 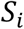, the outer loop takes the adapted parameters 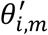, obtained in the inner loop, and evaluates the loss on the query set of task *m*. The resulting post-adaptation query-set losses from all tasks in *S*_*i*_ are then backpropagated to update the meta-parameters from the previous meta-training step, *θ*_*i–1*_ using *meta-learning rate β*. The updated parameters, denoted *θ*_*i*_ serve as input to the inner loop of the next meta-training step. After the final training step *N,θ*_*N*_ are the parameters of the output *pretrained model*. The pretrained model is, therefore, an initialization that minimizes the post-adaptation query-set loss across a distribution of tasks, aligning with the central objective of meta-learning: to learn model parameters that facilitate effective adaptation to new tasks.

Formally, the meta-parameters were updated according to the following update rule 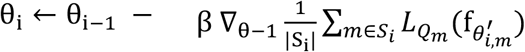 (Eq. 1). Here, *θ*_*i–1*_ denotes the meta-parameters at the beginning of meta-training step *i, S*_*i*_ is the mini-batch of meta-training tasks sampled at that step, |S_i_| is the number of tasks in the mini-batch, 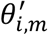denotes the parameters after inner-loop adaptation on task 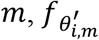is the adapted model for task 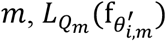 is the loss of the adapted model on the query set of task *m* (*Q*_*m*_), ∇_*θ*–1_ is the gradient of the average query-set loss (across entire mini-batch) with respect to the pre-adaptation meta-parameters, and *β* is the outer-loop learning rate.

The hyperparameters for the MAML-TabNet framework were empirically tuned to optimize loss on the meta-validation set of tasks.

We tracked performance along the meta-training steps. For each task, the model was first adapted on the support set and then the loss was evaluated on the query set. Analysis of meta-training tasks: at the end of every meta-training step *i*, the TabNet model with *θ*_*i*_ parameters was evaluated on all tasks from the mini-batch. 2) Analysis of meta-validation tasks: at the end of every epoch (namely, at the end of every 9 meta-training steps), the TabNet model was evaluated on 10 randomly-sampled (without replacement) meta-validation tasks. The loss was averaged over 100 separate initializations on 10 tasks randomly sampled. We further quantified the Δloss metric as adaptation-induced changes in each meta-training step.

### Data curation and task formulation

We curated a large collection of bulk RNA-Seq datasets downloaded from GEO, focusing on human studies. Overall, 2,079 datasets were used. To reduce the dimensionality of the input space, we selected the top 1,000 most variable genes across all datasets. The full collection of 2,079 datasets was partitioned into a meta-training set (*D*_*train*_, 1,479 datasets), a meta-validation set (*D*_*val*_, 300 datasets), and a meta-testing set (*D*_*test*_, 300 datasets), with no dataset overlap between splits. We focused on binary classification tasks, each defined within a specific dataset. Trivial meta-testing tasks, defined as tasks for which 1-nearest-neighbor classification achieved perfect accuracy, were excluded from all performance evaluations (423 tasks were excluded). Each meta-testing task was annotated using multiple classification schemes, including annotations by research objective, disease or cell population^22^.

### Performance evaluation

We compared GGE’s performance against *K* nearest neighbors (NN-*K*), SVM, decision tree (DT), random forest (RF), AdaBoost, XGBoost, and MLP. In addition, to evaluate the contribution of meta-training, we compared GGE against its standalone base learner (TabNet) and a Prototypical Network (ProtoNet) utilizing the same TabNet base learner. The same evaluation approach was applied to both GGE and each compared method: for each meta-testing task separately, the model was first adapted on the support set and then assessed on the query set. Performance for each task is computed as the average of 100 independent runs, each initialized with a distinct random seed. We evaluated performance using F1 score, precision, recall, and accuracy. Standard classifiers were implemented using Python’s scikit-learn and XGBoost libraries, while ProtoNet and TabNet were implemented using PyTorch and pytorch-tabnet. ProtoNet is based on code adapted from https://github.com/cnguyen10/few_shot_meta_learning.

The accuracy gains are most pronounced between 1-shot and 5-shot classifiers, after which the learning curves of most compared models effectively plateau, showing minimal marginal improvement with additional samples (data not shown). Based on these trends, we selected the 5-shot classifiers (*k*=5) as the primary setting for this study.

### Feature importance and pathway enrichment analysis

A primary advantage of using TabNet as the base learner is its inherent ability to provide feature explainability based on the weights of the attention masks^19^. To identify the specific genes that were used in each decision step of each task, we leveraged this property by calculating feature importance scores that utilize the sparse attention masks of the task-specific adapted model. For each task, the attention weights of the adapted network were extracted for the query set samples and aggregated across decision steps to obtain gene-specific importance scores. These scores were then averaged across all samples within a given task. A *global importance score* was calculated for each gene by averaging its importance scores across all meta-testing tasks. In addition, importance scores for each disease category were calculated by averaging each gene’s scores across the tasks belonging to each disease category.

We first selected the 100 genes with the highest global importance scores and tested their enrichment in each GO biological process (GO-BP) gene set using a hypergeometric test. This analysis identified 65 enriched gene sets (FDR *q* < 0.05). We then repeated the enrichment analysis for these 65 gene sets using only the 1,000 feature genes as the background, yielding 41 significantly enriched gene sets (*q* < 0.05). Finally, we tested the enrichment of each of these 41 gene sets using the disease-category-specific gene-importance scores. Throughout this analysis we used the GGE classifier with the best average F1 score across the validation tasks.

### Implementation

GGE is implemented in Python (v3.9) using PyTorch and pytorch-tabnet (TabNet). MAML is based on code from https://github.com/cnguyen10/few_shot_meta_learning.

### Data Availability

No new data were generated as part of this study.

## RESULTS

### Construction and validity of the GGE model

Several lines of evidence support the validity of our GGE model. First, we assessed model improvement during meta-training by tracking loss in each meta-training step. For each task, the model was first adapted on the support samples and then evaluated on the query samples, and scores were aggregated across tasks. The model showed consistent improvement on both meta-training and meta-validation tasks (**Figure 1E**).

Second, we quantified the benefit of task-specific adaptation at each training step by comparing query-set performance before and after adaptation. We summarized these adaptation-induced changes using the Δloss metric (implemented as slope; negative Δloss indicates improvement; evaluated on the meta-validation set). We observed that Δloss improves over meta-training steps (**Figure 1F**), indicating progressively more effective few-shot adaptation. Based on this analysis, we selected the parameters saved after 41 meta-training steps (*θ*_41_) as the pretrained model. Due to the nested-loop architecture of MAML, these 41 steps correspond to more than 295,000 inner-loop gradient updates (41 meta-steps ×120 tasks × 60 inner gradient steps). Although loss improved slightly in later steps, inter-task variance widened (**Figure 1E**). We therefore applied early stopping at meta-training step 41 to prioritize consistent generalization across diverse tasks.

Finally, GGE was compared with alternative methods. We observed that GGE’s performance correlates with the performance achieved by decision tree, random forest, and AdaBoost, all of which are decision-tree–based models (data not shown). This concordance is consistent with the design of GGE’s base learner (TabNet), whose sequential attention mechanism performs instance-wise feature selection and captures hierarchical feature interactions analogously to recursive splitting in decision trees.

### GGE performs well across diverse research objectives

Across the meta-testing tasks, GGE outperforms all alternative methods. GGE achieved a mean F1 score of 0.61, compared to F1 scores below 0.52 for all competing methods (**Figure 2A, left**). Furthermore, the distribution of task-level F1 scores showed a substantial shift in favor of GGE (paired t-test *q* <10^-17^; **Figure 2AB**).

**Figure 2.**
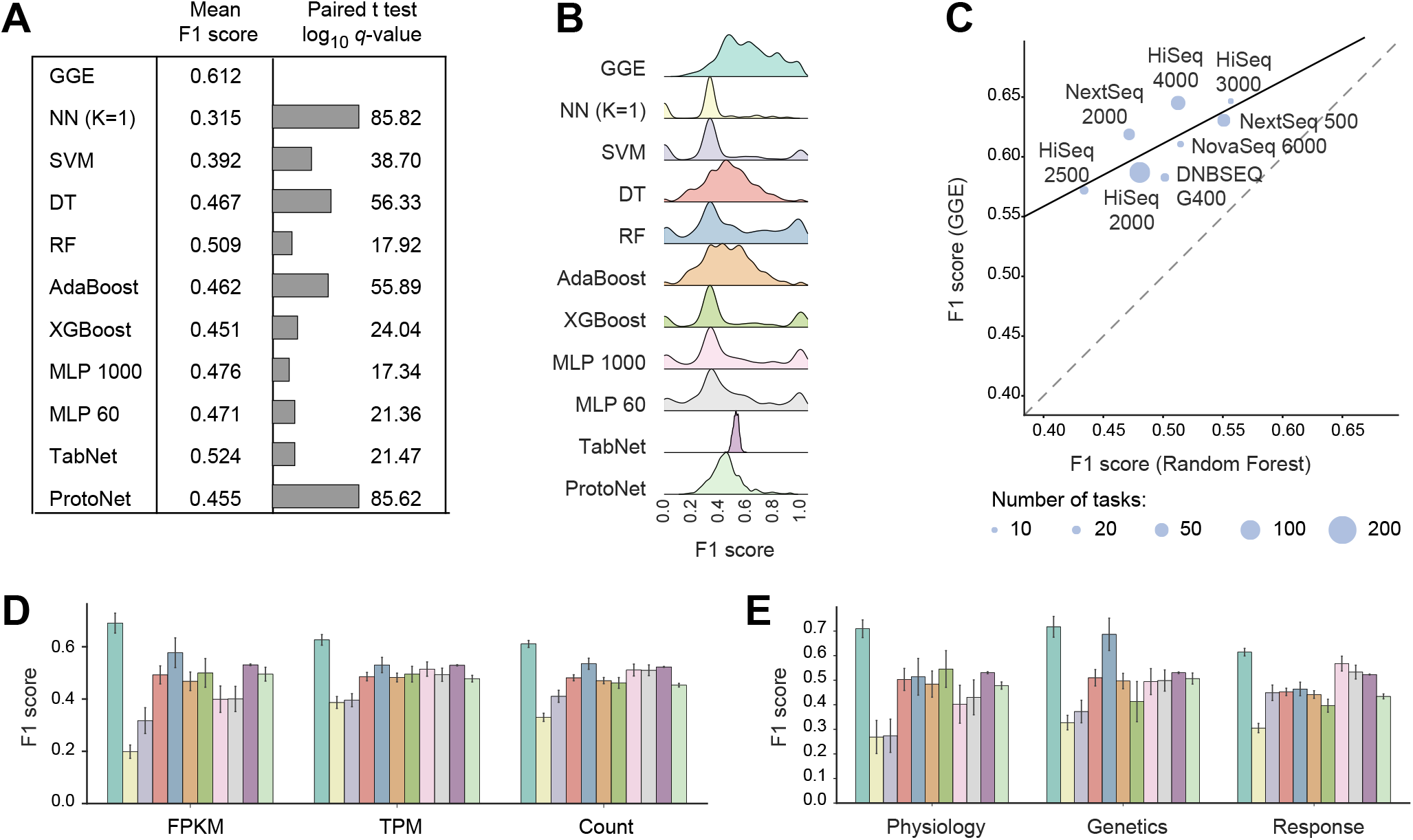
GGE outperforms alternative classifiers across diverse human diseases. Results are reported across meta-testing tasks. (**A–B**) Comparison of GGE with the alternative methods using mean F1 scores (A, left), pairwise comparisons between GGE and each compared method using paired *t*-test > values (A, right), and distributions of task-level F1 scores (B). The panels show that GGE is shifted toward higher F1 scores, whereas the compared methods exhibit either bimodal distributions or unimodal distributions centered at 0.5 or lower. **(C)** Performance across RNA-seq platforms. F1 scores are compared between GGE and random forest, with each point representing the mean performance across all tasks from a given RNA-seq platform. Only platforms represented by at least 10 tasks were included. **(D)** Performance across the most common RNA-seq preprocessing methods: FPKM, TPM, and raw gene counts (*n* = 49, 144, and 97 tasks, respectively). (**E**) F1 scores are shown for GGE and the compared methods across three broad biological domains. In D-E, bars: mean F1 score; error bars: standard error across tasks.

The meta-testing tasks were collected from multiple independent studies that differed in RNA-seq platform and preprocessing method, including TPM and FPKM. Two lines of evidence indicate that this technical heterogeneity did not drive the observed performance differences. First, GGE maintained strong performance across all RNA-seq platforms (**Figure 2C**). Particularly, platform-level GGE performance is correlated with random-forest performance (where each task is classified independently; **Figure 2C**), suggesting that the apparent platform-associated differences largely reflect variation in task difficulty rather than platform-specific effects. Second, we used only per-dataset scaling, without explicit batch correction or cross-dataset harmonization, allowing GGE to learn directly from this technical heterogeneity. GGE maintains strong performance across datasets processed using different methods (Fisher-combined paired *t*-test, *P* < 0.005; **Figure 2D**), supporting its robustness to preprocessing variation. Therefore, all evaluations were performed jointly across tasks from all platforms.

To further assess the advantage of GGE across diverse research objective, we annotated the meta-testing tasks using three domains: (i) *physiology*, encompassing comparisons of physiological states such as healthy vs. disease, disease subtypes (e.g., severe vs. moderate severity) (16 tasks);(ii) *response*, capturing the effect of intervention (treated vs. placebo or untreated samples), including both *in vivo* and *in vitro* interventions (30 tasks); and (iii) *genetics*, comprising comparisons between genotypes, including natural variation and experimental genetic perturbations (130 tasks). Across all three domains, GGE achieved higher F1 scores than all compared methods (**Figure 2E**). The mean F1 scores for GGE were 0.7 (physiology), 0.61 (response), and 0.71 (genetics), whereas the compared methods achieved mean F1 scores below 0.54, 0.56, and 0.68, respectively. These improvements were significant across methods – for example, GGE outperformed random forest (Fisher-combined paired t-test 9 < 10^−10^, a domain-level comparison, aggregated across domains). Collectively, these results indicate that GGE enables effective few-shot classification across diverse research objectives.

### GGE performs well across diverse disease types

To assess the relevance of GGE across diseases, we annotated each task with its corresponding disease category using the main chapters of the ICD-10 annotation (International Classification of Diseases, 10^th^ Revision; **Methods**). We identified six disease categories with sufficient representation (≥4 transcriptomic datasets and ≥4 tasks; **Figure 3A**). For each disease category, we summarized performance by averaging per-task F1 scores across all tasks assigned to that category. GGE achieved higher mean F1 scores than all compared methods in five of the six categories (**Figure 3A**). Moreover, GGE significantly outperformed each compared method across disease categories (Paired Wilcoxon test *q* <0.05 for all compared methods; **Figure 3B**), indicating robust performance across a broad range of diseases.

**Figure 3.**
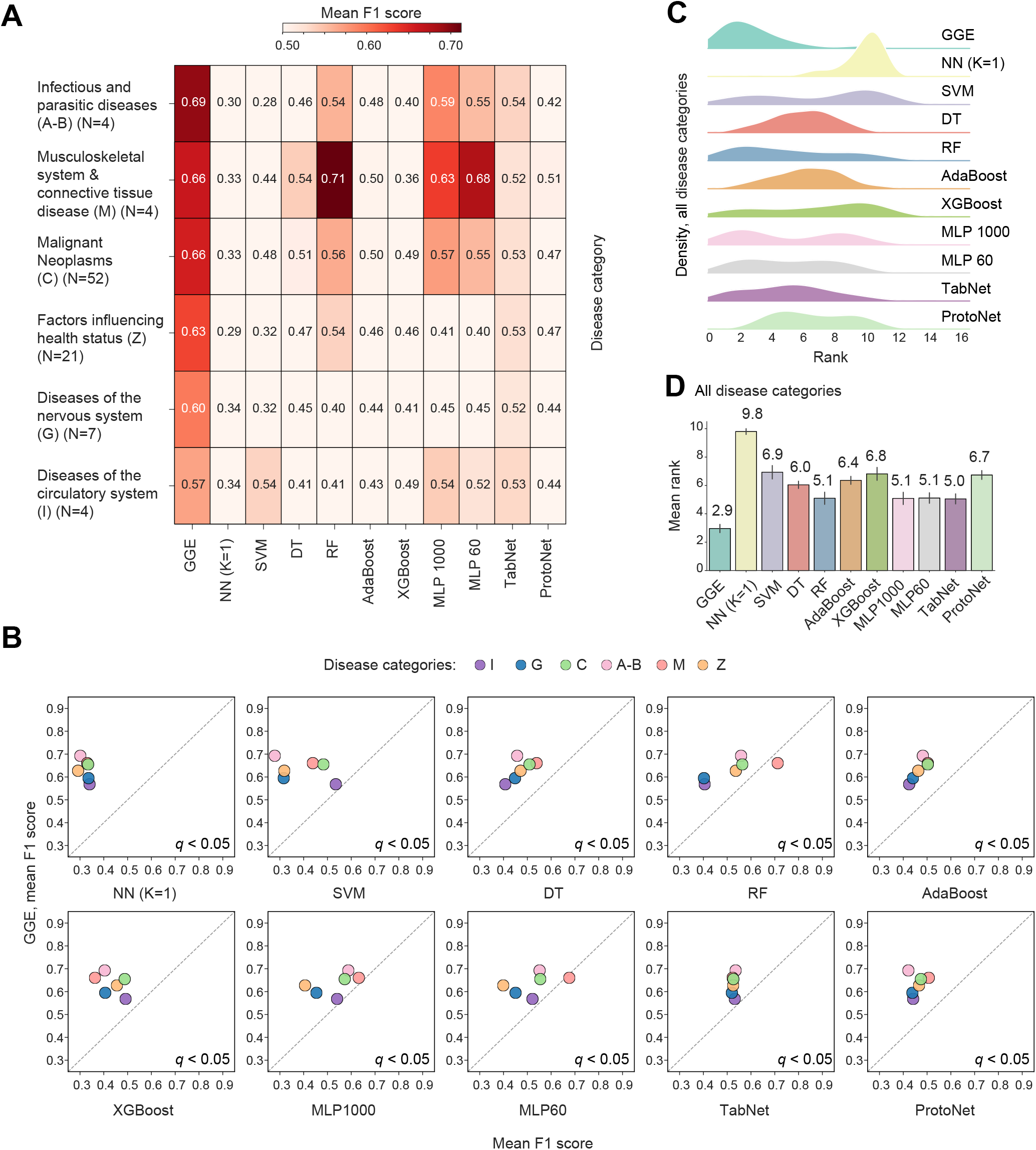
GGE performs robustly across diverse human diseases. Performance is shown across major ICD-10 disease categories (A,B) and at a finer clinical resolution across specific ICD-10 codes (C–D). Included are categories and codes with sufficient data (categories: ≥4 datasets, ≥4 tasks; codes: ≥1 task). N: number of meta-testing tasks. A,M,C,Z,G,I: ICD-10 categories. (**A–B**) F1 scores are shown as a matrix across disease categories and compared methods (A), and as scatter plots comparing GGE with each alternative method (B). (**C**,**D**) Ranking of methods by F1 scores. Within each code, classifiers were ranked by mean F1 score, with lower ranks indicating better performance. Ridge plots show the rank distribution of each method across the ICD-10 codes(C) and the bar plot shows the mean (and standard error) rank across the ICD-10 codes (D). GGE achieved significantly lower (better) ranks than all other methods (paired Wilcoxon test, *q* < 0.003).

We next evaluated performance at finer resolution using ICD-10 disease codes that correspond to specific diagnoses within each disease category (**Methods**). Across the meta-testing tasks, we identified 57 associated ICD-10 codes. Among these, 47 codes (82%) achieved a mean F1 score greater than 0.5, with an overall mean F1 score of 0.65 across codes. To systematically compare methods at the level of individual diagnoses, we ranked classifiers within each ICD-10 code based on their mean F1 score across tasks assigned to that code (lower ranks indicate better performance). This produced, for each method, a distribution of ranks across ICD-10 codes (**Figure 3C**). GGE achieved significantly better (lower) ranks than each compared method (paired Wilcoxon test *q* < 0.003; **Figures 3C,D**). For example, GGE attained the lowest mean rank (2.9), outperforming the next-best alternatives (TabNet, random forest, and MLP with hyperparameters of 60 and 1000 gradient steps), which had mean ranks between 5.0 and 5.1 (**Figure 3D**).

### GGE performs well across multiple cell types

To examine performance across diverse cell types, we used the DepMap collection^22^ as a standardized vocabulary of cell lines. We assigned each meta-testing task to a DepMap cell line (when relevant), yielding 19 cell lines with at least one task. For each cell line, we summarized performance by averaging task-level performance across all tasks assigned to that cell line. Across these 19 cell lines, GGE achieved robust performance: 95% of cell lines (18/19) had a mean F1 score > 0.5, with an overall mean F1 score of 0.7 across cell lines (**Figure 4A**). To compare methods systematically at the level of cell lines, we ranked classifiers within each cell line by their mean F1 score across tasks (lower rank indicates better performance), generating a distribution of ranks across cell lines for each method (**Figures 4B,C**). GGE achieved significantly better (lower) ranks than each of the compared classifiers (paired Wilcoxon test *q* <0.006; **Figure 4B**). Moreover, GGE attained the best mean rank (2.27), substantially outperforming the next-best alternatives (TabNet), which achieved a mean rank of 4.64 (**Figure 4B**).

**Figure 4.**
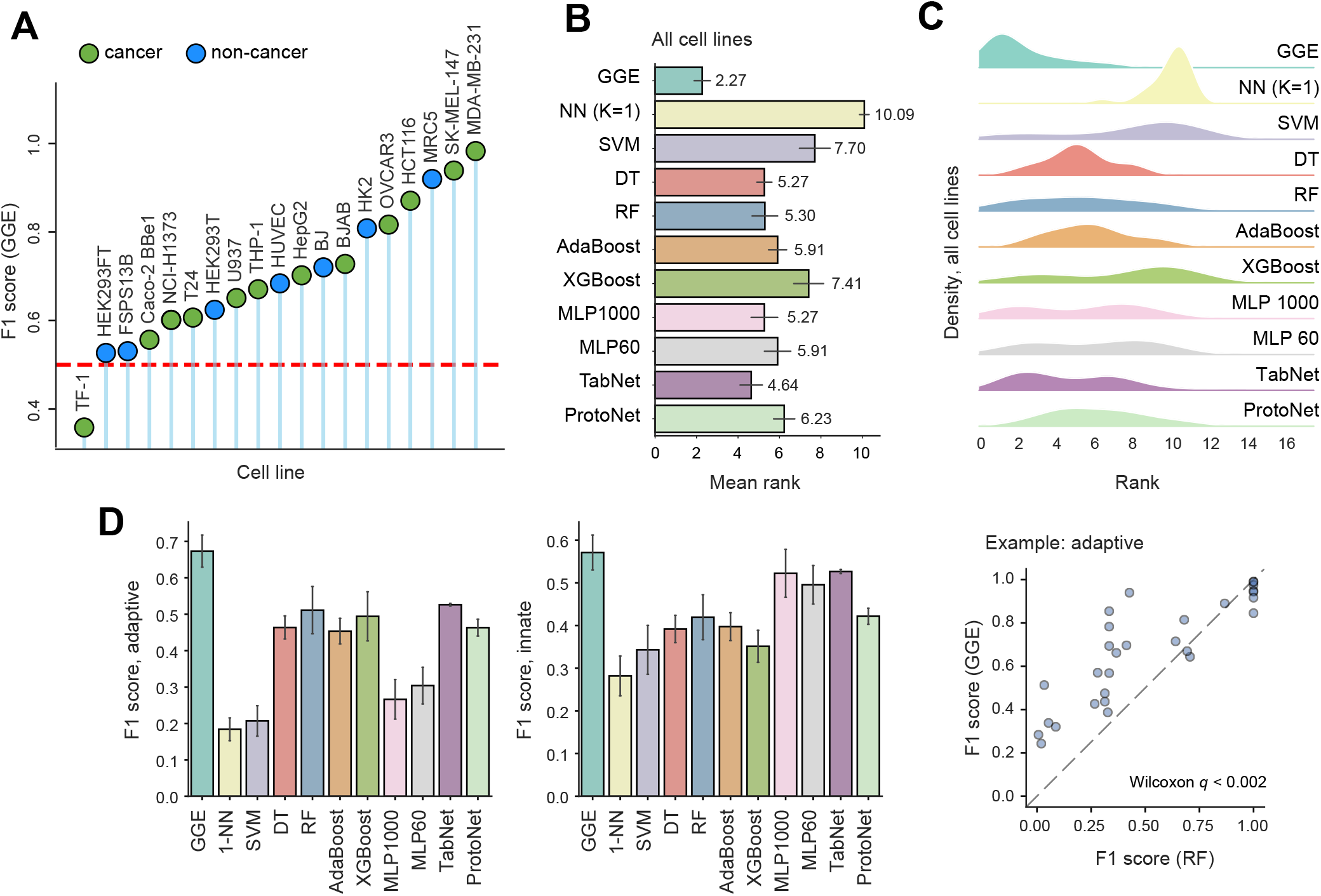
GGE performance across cell populations. (**A–C**) Cell line analysis. (**A**) Mean F1 scores across cell-lines. Color indicates whether the cell line originated from a cancer sample. (**B,C**) The rank of GGE and the compared methods across cell lines. (**D**) Classification performance across tasks involving adaptive (left) and innate (middle) immune cell populations. Right, representative comparison of F1 scores for GGE versus random forest across adaptive immune-cell tasks. Each point represents a distinct classification task. Dashed lines indicate the identity line.

Next, we analyzed performance across tasks annotated with either innate or adaptive immune cell types (22 and 29 tasks, respectively; **Methods**). GGE achieved the highest mean F1 score in both innate and adaptive populations (**Figure 4D,E**): 0.67 in adaptive immune populations and 0.57 in innate immune populations, whereas all comparison methods scored below 0.53 in both cell populations. This advantage was significant (paired Wilcoxon test, *q*-value < 0.05) for GGE against all methods in adaptive immune populations and for 7 of 10 methods in innate immune populations.

### Shared determinants of performance across biological objectives

Because GGE performed well across a wide range of biological objectives, we next asked whether common genes contribute to classification performance across multiple task types. Using attention-based analysis, we assigned each gene an importance score within each task and then averaged these scores across all meta-testing tasks. This yielded a *global importance score* for each gene, reflecting its recurrent contribution to classification across biological contexts (**Methods**). The top 100 genes with the highest global importance scores were enriched for multiple biological processes (41 enriched gene sets from the GO’s biological process collection, FDR *q* <0.05). These processes were related to core cellular functions and maintenance of homeostasis, including membrane transport (e.g., genes SLC17A5, PKD2, CHRNA10, and CLIC1), lipid metabolism (e.g., genes FASN, GPAT4, and DECR1), and vesicle mediated transport (e.g., genes VPS54, GOLPH3, COG2, and GGA2). Thus, genes that recurrently contribute to performance across diverse biological objectives appear to reflect fundamental cellular processes whose dysregulation is relevant across multiple conditions.

We then asked whether these pathways showed consistent contributions across disease categories. To this end, we repeated the same attention-based analysis separately within each disease category (same disease categories as in **Figure 3A**). Among the 41 significantly enriched processes identified in the global analysis, 37 processes were enriched at FDR *q* < 0.05 in at least five of the six disease categories (exemplified in **Table 1**). Overall, these results suggest that the shared genes used by GGE reflect core biological functions that contribute to classification across diverse disease contexts.

**Table 1.** Global feature importance across biological objectives. The table reports enrichment based on global gene-importance scores (column 1) and disease-category-specific importance scores (columns 2–7). All analyses were performed using meta-testing tasks, and all values represent FDR-adjusted *q* values.

| Gene set | Global *q*-value | Infectious and parasitic diseases (A-B) | Diseases of the circulatory system (I) | Musculo-skeletal system & connective tissue (M) | Diseases of the nervous system (G) | Factors influencing health status (Z) | Malignant Neoplasms (C) |
| --- | --- | --- | --- | --- | --- | --- | --- |
| Transmembrane transport | 0.001 | 0.000 | 0.004 | 0.055 | 0.002 | 0.002 | 0.004 |
| Electron transport chain (ETC) | 0.009 | 0.009 | 0.009 | 0.089 | 0.009 | 0.012 | 0.010 |
| Ameboidal type cell migration | 0.017 | 0.017 | 0.045 | 0.065 | 0.019 | 0.022 | 0.021 |
| Cellular response to ketone | 0.020 | 0.020 | 0.024 | 0.086 | 0.021 | 0.010 | 0.023 |
| Regulation of glucose import | 0.021 | 0.022 | 0.023 | 0.058 | 0.023 | 0.029 | 0.024 |
| Lipid metabolic process | 0.023 | 0.012 | 0.024 | 0.141 | 0.024 | 0.037 | 0.064 |
| Response to oxygen-containing compound | 0.026 | 0.026 | 0.028 | 0.224 | 0.027 | 0.018 | 0.021 |
| Regulation of spindle microtubules | 0.028 | 0.030 | 0.032 | 0.047 | 0.032 | 0.035 | 0.033 |
| Vesicle mediated transport | 0.030 | 0.047 | 0.047 | 0.058 | 0.047 | 0.037 | 0.033 |
| Cell adhesion | 0.029 | 0.013 | 0.022 | 0.057 | 0.030 | 0.025 | 0.062 |

## DISCUSSION

Our work provides, for the first time, evidence that the integration of multiple datasets can enhance the computational analysis of new datasets, without relying on any prior knowledge. We constructed a MAML–TabNet model and meta-trained it on thousands of binary transcriptomic classification tasks systematically derived from public human bulk RNA-seq studies. The resulting GGE model is designed to provide a reusable initialization for few-shot learning in human transcriptome data. Using held-out datasets, GGE improves performance relative to conventional classifiers and the established few-shot baseline, ProtoNet. This advantage is observed across a wide variety of tasks, including various research objectives, disease categories, and isolated cell populations, supporting the use of a single learned initialization as a broadly applicable starting point for transcriptome-based classification in low-sample settings. GGE also maintained strong performance across RNA-seq platforms and preprocessing pipelines, indicating robustness to technical heterogeneity. Notably, the utility of GGE for tasks derived from isolated cell populations argues that it captures cell-intrinsic signals, rather than relying primarily on differences in cell-type composition.

A key open challenge for meta-learning from public transcriptomic resources is representation bias in the meta-training and meta-validation tasks. Although datasets were selected from GEO without favoring particular diseases, experimental systems, or research questions, the repository itself may be inherently biased because some biological contexts are studied much more extensively than others. Moreover, there is no established “correct” distribution of diseases, pathways, biological systems, or experimental designs against which such biases could be identified or corrected. Encouragingly, we found no evidence that greater representation during meta-training conferred a performance advantage. *Response* tasks comprised 27.7% of meta-training tasks yet showed lower meta-test performance than the less frequent *genetics* (9.5%) and *physiology* (3.7%) domains. Similarly, disease categories that were more prevalent during meta-training did not systematically achieve higher meta-test performance. Thus, the major representation imbalances examined here do not appear to dominate the learned GGE initialization. Nevertheless, subtler biases may remain, and developing principled strategies for constructing biologically representative meta-learning collections is an important direction for future work.

A key insight from our analysis is that both components of the framework – meta-training and task-specific adaptation make substantial and complementary contributions to performance. First, comparison of GGE with a standalone base learner (TabNet) isolates the value of the meta-trained initialization. The two models use the same TabNet architecture, the same classifier hyperparameters, and the same training procedure on each new task, differing only in their initialization: GGE uses the initialization learned through meta-training, whereas standalone TabNet starts from random initialization. The improved performance of GGE over standalone TabNet therefore directly demonstrates the benefit of the meta-trained model initialization. Second, Δloss directly quantifies the contribution of task-specific adaptation, because it compares performance before and after adaptation to a new task. In our data, Δloss is consistently improved, indicating that the adaptation step adds value beyond that of the pretrained model obtained via meta-training. Together, these results show that GGE benefits from both the general initialization and effective adaptation to new tasks, underscoring the complementary roles of meta-learned priors and task-specific adaptation.

## CONCLUSIONS

In this study, we demonstrate improved classification based on a limited number of transcriptomic samples through the application of our General Gene Expression (GGE) framework. By applying meta-training across thousands of heterogeneous public datasets, GGE learns a robust initialization that improves generalization across diverse biomedical domains. By generalizing across biological objectives, diseases, and cell types, without requiring closely matched source data, GGE maximizes the utility of public transcriptomic repositories. This provides a general-purpose, data-efficient foundation to accelerate biological and clinical discovery in data-scarce settings.

